# Multiple Binding Configurations of Fis Protein Pairs on DNA: Facilitated Dissociation versus Cooperative Dissociation

**DOI:** 10.1101/696054

**Authors:** Min-Yeh Tsai, Weihua Zheng, Mingchen Chen, Peter G. Wolynes

## Abstract

As a master transcription regulator, Fis protein influences over two hundred genes of *E-coli.* Fis protein’s non-specific binding to DNA is widely acknowledged, and its kinetics of dissociation from DNA is strongly influenced by its surroundings: the dissociation rate increases as the concentration of Fis protein in the solution-phase increases. In this study, we use computational methods to explore the global binding energy landscape of the Fis1:Fis2:DNA ternary complex. The complex contains a binary-Fis molecular dyad whose formation relies on complex structural rearrangements. The simulations allow us to distinguish several different pathways for the dissociation of the protein from DNA with different functional outcomes, and involving different protein stoichiometries: 1. Simple exchange of proteins and 2. Cooperative unbinding of two Fis proteins to yield bare DNA. In the case of exchange, the protein on the DNA is replaced by solution-phase protein through competition for DNA binding sites. This process seen in fluorescence imaging experiments has been called facilitated dissociation. In the latter case of cooperative unbinding of pairs, two neighboring Fis proteins on DNA form a unique binary-Fis configuration via protein-protein interactions, which in turn leads to the co-dissociation of both molecules simultaneously, a process akin to the “molecular stripping” seen in the NFκB/IκB genetic broadcasting system. This simulation shows that the existence of multiple binding configurations of transcription factors can have a significant impact on the kinetics and outcome of transcription factor dissociation from DNA, with important implications for the systems biology of gene regulation by Fis.

## Introduction

Protein-DNA assemblies fundamentally are molecular machines that control the flow of genetic information to fulfill cellular functions.^1^ Dynamic protein-DNA interactions influence how DNA-binding proteins search and find their target sites along DNA and also how these proteins fall off the DNA, thereby regulating the expression of genes near these binding sites. In the simplest models, the concentration dependent thermodynamics of the binding and unbinding of proteins on DNA acts as a molecular switch for activating or deactivating the gene.^2^ Recently, a number of studies have pointed out that protein concentration not only influence the binding process through mass action but also surprisingly can catalyze protein dissociation (unbinding) from DNA in an unexpected way, presumably due to the multivalent nature of many transcription factor binding interactions (see a recent review by Chen et al.).^3^ Recent studies on the concentration-dependent protein dissociation (or facilitated dissociation) have opened a door to understanding the kinetic aspects of gene regulation.

Fis protein is involved in a variety of functions in *E-coli.* Fis provides mechanical support during DNA recombination;^4^ It also functions as a transcription factor and regulates numerous genes;^5^ It also bends DNA in order to assist chromosome packaging. It is surprising that a protein with such an apparently simple dimeric topology can demonstrate this great diversity of functions. Fis protein shows an interesting dissociative behavior arising from its dimeric nature. Recent experiments show that the turnover rate of fluorescently labeled bound Fis protein on DNA is coupled to the concentration of Fis in the surrounding solution.^6^ This unexpected concentration dependence arises from a process that has been called facilitated dissociation. Facilitated dissociation processes seem to occur for proteins having multiple domains. Multivalent binding allows a protein to become partially dissociated so that some partial protein-DNA contacts are broken through thermal fluctuations. One plausible mechanism for the concentration dependent kinetics thus involves the formation of a ternary complex where a second protein from solution competes with the already bound one for the same or a nearby DNA-binding site, thereby favoring a partially dissociated intermediate.^6–11^ This competition effect was uncovered by Kamar et al. at the single-binding site level using single-molecule imaging.^10^ Facilitated dissociation shows how nature can exploit a simple binding-site competition strategy that relies on the multi-meric binding of proteins to carry out nonlinear logical responses to environmental stimuli. In living cells, the regulation is not always quasi thermodynamic but can be kinetically controlled. Presumably, to achieve kinetic control requires accelerating the dissociation through high-affinity interactions with binding competitors. Under such control, the residence time of receptor-ligand complexes rather than their binding equilibrium becomes the most significant factor for biological function.^12^ Catalysis of dissociation can change the residence time of transcription factors on DNA and thus affect their regulatory functions and especially influence the stochastic aspects of gene expression.^13,14^ Facilitated dissociation has also been observed in bacterial chromosome assembled *in vivo*^15^ suggesting that this phenomenon is a general biological phenomenon in living cells.

Quantitative treatment of protein-DNA binding is often based on a dynamic scheme of protein association with and dissociation from DNA, that at equilibrium would be quantified by the thermodynamic dissociation constant *K*_d_ = *k*_off_ / *k*_on_. Often in systems biology, the backward rate coefficients are assumed to be concentration-independent and would reflect only spontaneous dissociation. Recently however, many experiments on several different systems have shown that *k*_off_ varies with the protein concentration in solution. The dissociation rates for the metalloregulator CueR,^16^ the bacterial protein HU,^6^ and the yeast HMGB protein NHP6A^6^ have all been found to be concentration-dependent. Fast polymerase turnover during DNA replication in the bacterial replisome also shows concentration dependence.^17^ Similar phenomena are also observed for replication protein A (RPA) dissociating from single-stranded DNA (ssDNA).^18^ Facilitated dissociation has also been seen for other metal-sensing transcription regulators in living cells.^19^ Here we show how energy landscape analysis and coarse-grained molecular dynamics simulations can give several insights into the physical mechanism of this unexpected behavior.

Previously, we have explained the concentration dependence of labeled Fis dissociation in terms of a three-state kinetic model, involving a partially dissociated configuration of the Fis:DNA complex.^8^ This intermediate takes part in a dynamic equilibrium with its bound Fis-DNA configuration. As a result, the population of the intermediate is controlled by the availability of other proteins in solution via a direct competition for a DNA binding site. This competition takes place between the protein molecule that has already been bound on DNA and the other proteins still in the solution phase. Our previous work highlighted one particular pathway for dissociation—facilitated dissociation.^8^ Nevertheless, we cannot exclude the influence of other possible kinetic routes involving still higher-order interactions within the protein-DNA assembly. One such proposal has already suggested in the literature.^7,10^ Sing et al. specifically put forward the idea that there are several different dissociation pathways resulting from there being a partially dissociated state. They divide the mechanisms into separate concentration-independent and concentration-dependent pathways.^7^ Their model does not take into account binding cooperativity of proteins on DNA. Another possibility is that the cooperative effects may come from protein-protein interactions on the DNA that changes the dissociation pathway. A similar kinetic scheme was recently adopted to account for the effect of salt on the protein dissociation rate.^10^ The molecular details underlying the cooperative interactions are still not clear. Here, we investigate these ideas by tracking the correlation of motion of multiple proteins on DNA. We specifically track how the protein-DNA complex is formed and examine from a molecular perspective how a partially dissociated Fis can interact with a partner Fis. Our results show there is a diverse ensemble of configurations of binary-Fis species on DNA (Fis1:DNA:Fis2 ternary complex). One particularly important conformation is a symmetric binary-Fis configuration where one of the two DNA-binding domains on each Fis molecule comes off from half of the binding site while the other DNA-binding domains remain bound to the binding site. In addition to this partially bound structural feature, there are significant physical contacts formed between the two Fis molecules, thereby forming a symmetric molecular construct on DNA. A similar ternary complex has been inferred experimentally for CueR metalloregulator using an engineered DNA.^20,21^ Chen et al.^21^ pointed out in a review article that the such a ternary complex (CueR:DNA:CueR) is likely a common intermediate for many different transcription deactivation pathways. Specifically, the incoming CueR can exchange for another one that is already bound; or the binary-CueR pair as a whole can fall off from DNA. The former outcome corresponds to facilitated dissociation that we have previously described^8^ while the latter outcome involves the idea of cooperative interaction that we focus on the present work. We show using computer simulations of realistic coarse-grained models that the binary-Fis configuration is indeed populated. This observation allows us to explore an alternative dissociation pathway, termed “cooperative dissociation,” where a solution phase Fis interacts with the Fis already on the DNA, forming a binary-Fis intermediate, having specific physical contacts between the two, prior to dissociation of both proteins from the DNA. Due to the stoichiometry change in cooperative dissociation, the outcome of this co-dissociation of both molecules has important implications in the systems biology of gene regulation by Fis.

In this work, we specifically study the structure, stability and cooperative binding of the binary-Fis intermediate. Here we generalize our previous analysis by incorporating various stoichiometries of protein and DNA and show that this theoretical framework is applicable to multiple protein-DNA assemblies. By exploring the binding energy landscape of Fis proteins binding a short piece of DNA, both of the dissociation scenarios are investigated, depending on the number of proteins being dissociated: *p* = 1 (facilitated dissociation) and *p* = 2 (cooperative dissociation), as illustrated in Fig. 1(a). We show that the existence of distinct binding configurations of pairs of Fis molecules on DNA leads to existence of distinct dissociation pathways. The formation of protein:DNA:protein ternary complexes seems to be a generic mechanism for modulating a range of biological functions in cells, including possible exchange of transcription factors and allows a nontrivial logic of transcriptional control.

**Figure 1.**
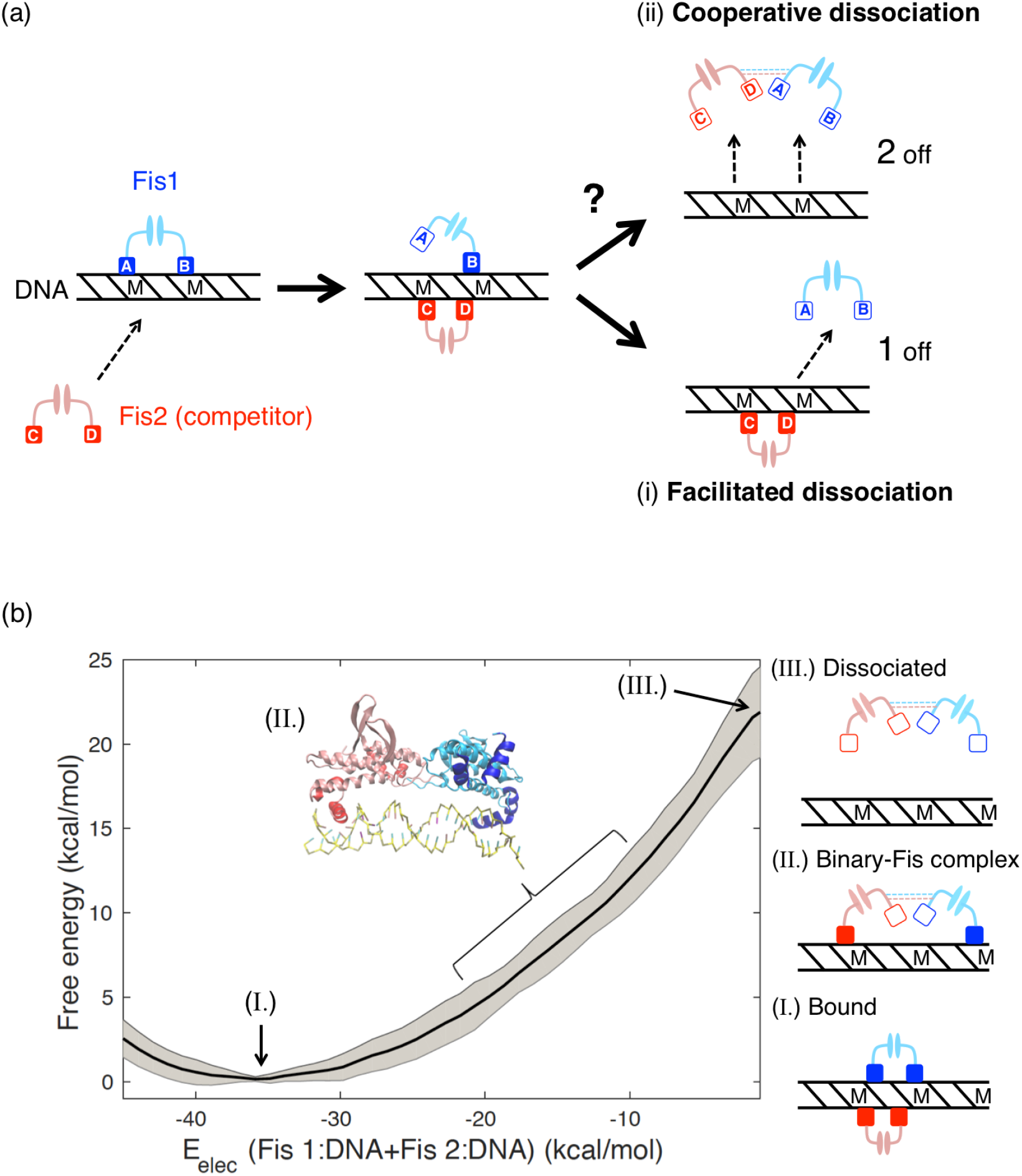
Distinct binding configurations of multiple Fis proteins are hypothesized to underlie different types of dissociation pathways, (a) A schematic diagram of multiple dissociation pathways. The red Fis protein (Fis2) competes with the blue Fis protein (Fis1) bound on the DNA for the binding site. Depending on the protein-DNA ternary configuration being visited, the proteins can undergo (i) facilitated dissociation or (ii) cooperative dissociation. The former leads to the net effect of protein exchange (only one protein comes off while the other one stays); the latter leads to an overall dissociation of the proteins together (both two proteins come off as a result). The arrows indicate the flow of the dissociation pathways. Note that for the cooperative dissociation channel we hypothesize the presence of a unique binary-Fis configuration prior to the overall dissociation, (b) The free energy profile as a function of the electrostatic interaction energy *E*_elec_ of the protein Fis1 (colored in blue) and Fis 2 (colored in red) with the DNA is shown. The binary-Fis configuration is observed in the sampling windows between *E*_elec_= −10 ~ −20 kcal/mol. Example configurations are shown as cartoon diagram on the right with labels I, II, III, which represent the bound, the binary-Fis, and the dissociation configurations, respectively. DNA is represented by a black strip with “M” letters showing the location of major grooves. Each Fis protein has two DNA-binding domains shown in square, and each domain can either bind (*solid*) or unbind (*hollow*) to the DNA. Physical contacts between the two Fis proteins are denoted by dotted lines.

## Results

### Discovering a new binary-Fis ternary configuration

We first explored the free energy profile of protein dissociation using the electrostatic interaction energy between the Fis protein and DNA as a reaction coordinate for tracking the dissociation process. To sample configurations, the total electrostatic energy of both proteins with DNA (Fis1:DNA+Fis2:DNA) was first used as a biasing coordinate in the weighted histogram method (WHAM). This choice of coordinate allows one to track simultaneously the configurations of both of the two Fis proteins on DNA along the dissociation path. Fig. 1(b) shows the free energy profile as a function of total electrostatic energy (*E*_elec_) of the interaction of the protein Fis 1 (colored in blue) and Fis 2 (colored in red) with the DNA. The bound state refers to a ternary complex where both Fis molecules are solidly bound to the DNA. This bound state is located at the global minimum (*E*_elec_ ≈ −35 kcal/mol) of the free energy profile. The negative sign and large value of the electrostatic interaction energy *E*_elec_ indicates there is a strong attraction between the Fis molecules and the DNA. The dissociated state refers to the situation where both Fis molecules are well separated from the DNA where *E*_elec_ ≈ 0 kcal/mol. One sees no significant intermediate well along this one-dimensional free energy profile (−35 < *E*_elec_ < 0 kcal/mol). In the simulation trajectories, however, one often finds on careful examination a binary-Fis ternary complex that is long-lived during the simulation. Although this binary-Fis complex does not seem to be well populated when sampled along the electrostatic biasing coordinate, this does not exclude the possibility that one can isolate a well defined population corresponding to it along other coordinates. Fig. 2 shows two example configurations of the binary-Fis ternary complex where each of the individual Fis proteins are partially bound to DNA but where they mutually make significant contacts with each other through protein-protein interactions. As a result, the two Fis proteins form a binary-Fis configuration on DNA. This configuration exhibits *C*_2_ chemical symmetry. It is worth noting that this binary-Fis structure as a whole can dynamically switch between two distinct orientational positions by sliding along the DNA. The sliding motion involves inserting one of the DNA-binding domains (the anchor) into the DNA major/minor grooves. The major grooves along the DNA form a wider helical channel that allows Fis proteins to walk along in a circular path. Consequently, the symmetric binary-Fis adduct can reposition itself through a sliding motion. Each position shows a specific orientation of the dimerized Fis proteins with respect to DNA. For example, if we draw a line between the two Fis molecules on the DNA, from a top view we would see this line forms an angle against the axial direction of the DNA. The change of the angle represents different orientations of the Fis pair with respect to the DNA. One orientation is parallel (0°) to the DNA, which we call the standing orientation (Fig.2(a)) while the other is perpendicular to the DNA which we call the riding orientation (Fig.2(b)). The symmetric binary-Fis configuration shown contains a molecular dyad (consisting of two Fis proteins). Using this symmetry, multiple Fis proteins can structurally rearrange on the DNA into a novel molecular construct through protein-protein interactions. Fig.2(c) shows the probability contact map for the basin corresponding to the symmetric binary-Fis configuration on DNA where the interprotein contacts are highlighted.

**Figure 2.**
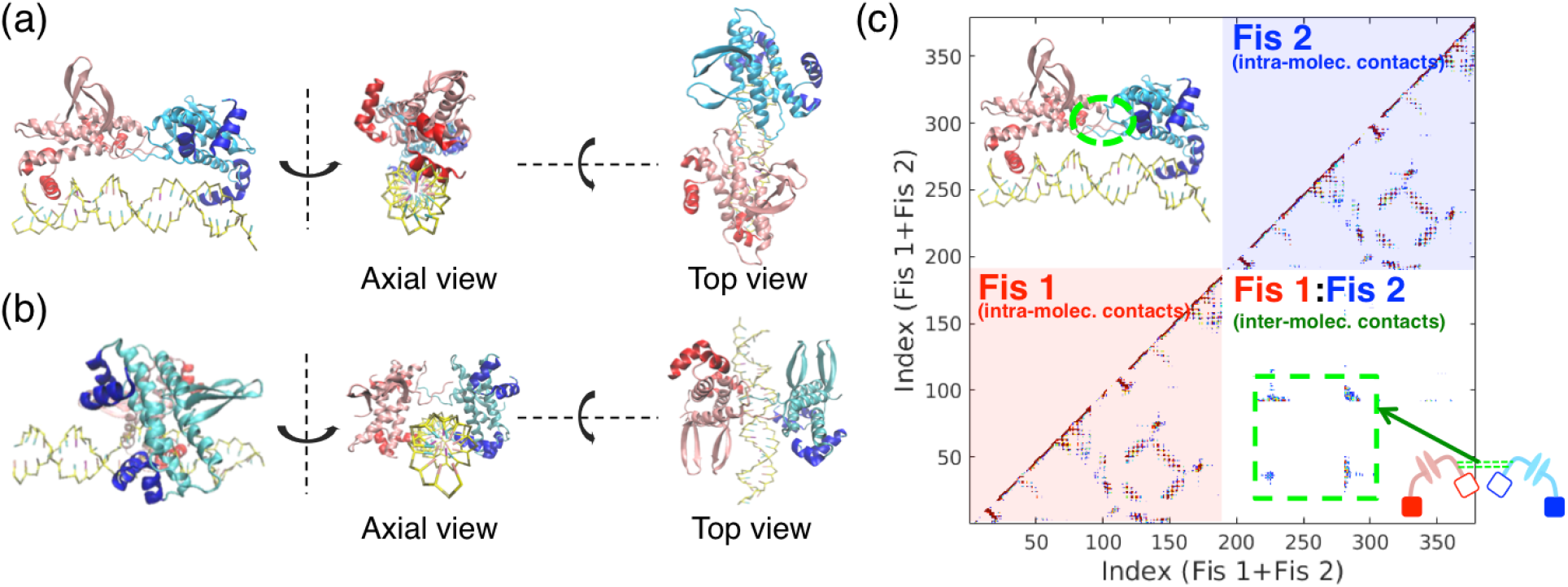
Examples of the binary-Fis protein-DNA configuration are shown, (a) The standing orientation, (b) The riding orientation. Both (a) and (b) show *C*_2_ chemical symmetry, (c) Probability contact map of the binary-Fis configuration.

We now examine the stability of the binary-Fis species. We can explore the population of binary-Fis by projecting the free energy surface onto additional coordinates beyond those used in the WHAM bias. In our previous study, we used the electrostatic interaction energy of one single Fis protein with the DNA *E_elec_*(Fis 1: DNA) as the progress coordinate for studying facilitated dissociation.^8^ Here we also explore other coordinates that are able to provide additional structural details about the binary-Fis configuration. To define these coordinates, we use a centroid structure taken from the ensemble of binary-Fis configurations as a reference structure and then calculate the *Q* value (*Q_binFis_*;, see Methods) for all snapshots from the simulation trajectories with respect to this fiducial structure. We also use the center-of-mass distance between Fis1 and Fis2 on the DNA (COM Distance) as well as the total number of protein contacts (*N_contact_*) as additional coordinates to separate more clearly distinct subpopulations of binary-Fis configurations. None of these intuitive progress coordinates turns out to be ideal (see Fig.S1). In the next section, we will show how better coordinates can be constructed using a dimension reduction technique, principal component analysis (PCA).

### Constructing free energy surfaces of the binary-Fis ternary complex using collective variable principal component analysis (PCA)

In order to analyze more clearly the binary-Fis configurations, we have employed collective variable (CV) principal component analyses to find optimal combinations of the intuitive trial collective basis variables that we introduced above. The input basis variables have clear meanings in terms of the various structural and dynamical features of the protein-DNA complex. Specifically, we use the following collective variables (CV1 to CV7) as inputs for a principal component analysis: 1. *E_elec_*(Fis 1:DNA+Fis2:DNA) 2. *E_elec_*(Fis 1:DNA) 3. *E_elec_*(Fis2:DNA) 4. the number of protein-protein contacts (*N_contact_*) 5. center-of-mass distance (COM distance) 6. the number of head-to-head protein contacts (*H_contact_*), and 7. the *Q* value of the binary-Fis configuration (*Q_binFis_*) with respect to a reference structure (see Methods). These collective variables are grouped and illustrated in Fig.S2. Items 1-3 describe the electrostatic interactions between the protein moieties (charged residues) and DNA phosphate groups; Items 4-5 explore general physical contacts and geometric position between proteins on DNA; Items 6-7 provide structural specificity that distinguishes several of the particular protein quaternary structures from each other. In contrast to looking at the free energy projected solely onto the single input trial collective variables, for which the profiles appear monotonic and featureless, using the collective variable PCA pulls apart the cross-correlations among these trial collective coordinates.

Fig. 3 shows several of the two-dimensional free energy surfaces obtained by plotting the free energy against various computed principal components (PCs). The collective variable principal component analysis uses 7 collective trial basis variables. Thus there are 7 new distinguished collective coordinates PC1 to PC7. These essentially represent a new basis set of transformed coordinates which are linear combinations of the original collective basis variables used as input to the PCA. The coefficients in these linear combinations are listed in Table 1 along with the percentage of total variance that is explained by each component. Each panel in the figure shows the free energy projected individually onto a single PC along with the common coordinate *E_elec_*(Fis 1:DNA). The first panel presents the free energy surface along PC1, which accounts for ~50% of total variance. PC1 is the most important PC. On the surface, the major basin corresponds to the bound state (*E_elec_*(Fis 1:DNA) = −20 kcal/mol) in which both Fis proteins are solidly bound. This bound major basin is also visualized clearly when employing different PCs as can be seen from panel to panel. One particular feature that emerges from looking at the PC1 surface is that there is a minor basin at *E_elec_*(Fis 1:DNA) = −10 kcal/mol. This basin does not show up clearly when using the other PCs. This result indicates a distinct population of binary-Fis configuration on the surface. Note that PC1 is composed of ~60% of *E_elec_* and ~15% of *Q_binFis_*(see Table 1). As the most important PC, we learn from this analysis that *Q_binFis_* provides enough structural specificity to help identify the population of the key binary-Fis configuration. To further the analysis for the binary-Fis populations, we proceeded to examine the rest of PC surfaces with significant fraction of *Q_binFis_* These are shown, for example, for PC3. PC3 is made of ~20% of *Q_binFis_*, ~20% of COM distance, and ~40% of *H_contact_*. In addition to *Q_binFis’_*, we find that *H_contact_* provides additional specificity such that when projecting the free energy onto both PC1 and PC3 on the surface, we can see that the symmetric binary-Fis basin is clearly separated from the major one (see Fig. 3(b)). The free energy difference between the binary-Fis basin (*F_binFis_*) and the major basin (*F* is set to be 0) therefore is estimated to be, Δ*F*_binFis_=4.2 kcal/mol. Here we show that multiple PCs allows us to identify the population of the binary-Fis configuration. The other PC surfaces are less informative, but they do provide insight into some other binary-Fis species, for example, PC2 and PC4 both explore configurations of generic binary-Fis ternary assemblies that form through non-specific protein contacts. These PC surfaces resemble those obtained by solely projecting over COM distance and *N_contact_* (see Fig.S1). The first four PCs (PC1 to PC4) account for over 90% of total variance of the data. The remaining PCs (PC5 to PC7) provide only <10% of the total variation. PC5 represents total electrostatics including *E_elec_*(Fis1:DNA+Fis2:DNA), *E_elec_*(Fis1:DNA). and *E_elec_*(Fis2:DNA): the free energy surface exhibits no clear substructure but rather one nearly featureless major basin in a linear shape. PC6 is somewhat interesting. Despite its extremely tiny contribution to the total variance 3.5%, PC6 embodies the highest degree of structural specificity including 50% of *Q_binFis_* and 30% of *H_contact_*. The corresponding free energy surface shows an intriguing pattern at *E_elec_*(Fis 1:DNA) = −10 kcal/mol with the major basin branching out into three different directions on the surface. This result suggests that multiple binary-Fis species undergo a dynamic interconversion equilibrium. PC7 is not of interest because it does not contribute to the total variance at all thereby the resulting profiles are featureless.

**Figure 3.**
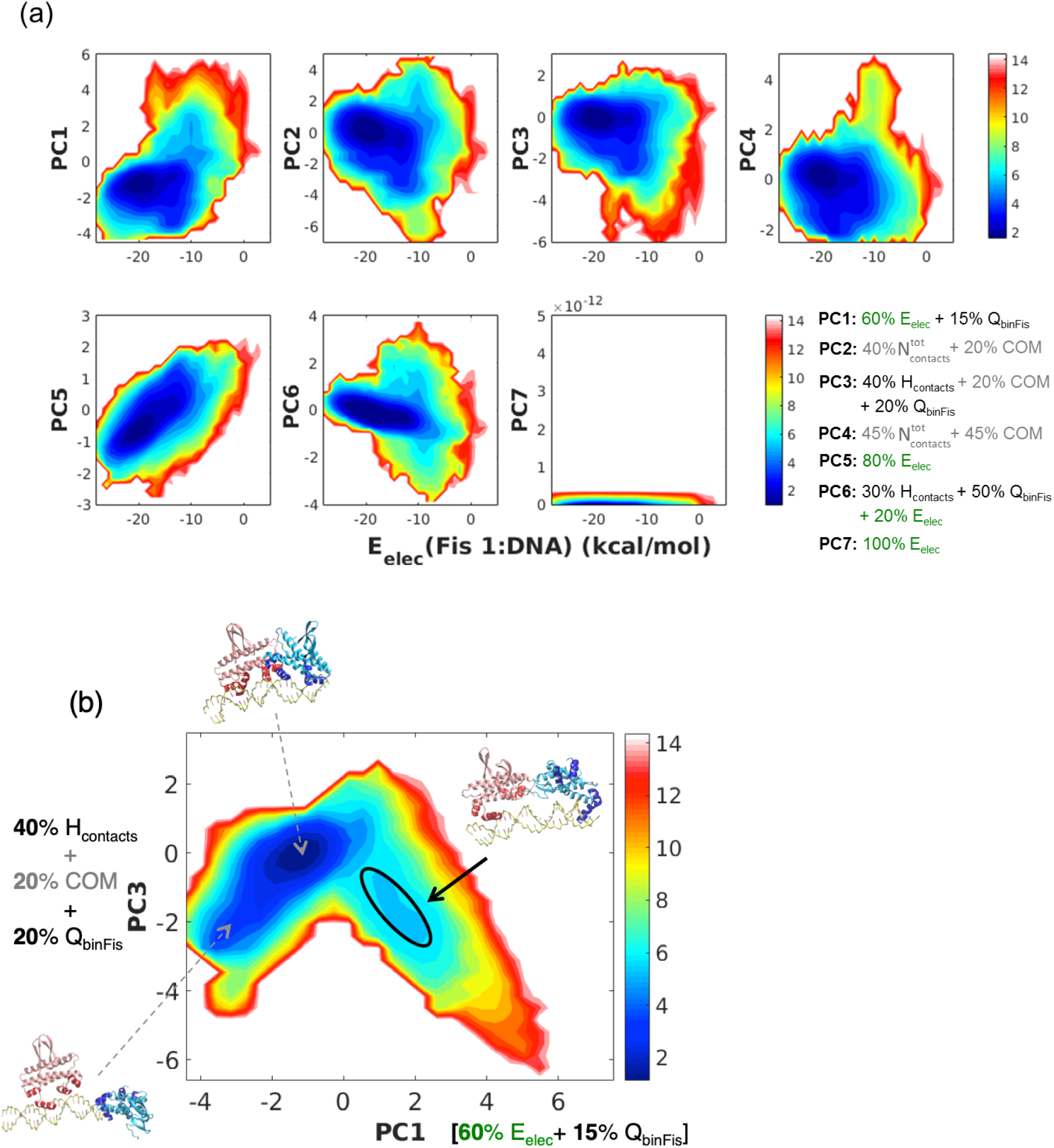
The mechanism of dissociation of Fis protein from DNA is explored using a two-dimensional free energy surface as a function of principal components (PCs), (a) Each free energy surface in the panels shows the free energy projected onto one particular PC (PC1 to PC7, sequentially aligned from left to right on top and then bottom) against *E*_elec_(Fis1:DNA). *E*_elec_(Fis1:DNA) spans from −24 to 0 kcal/mol with −20 indicating the bound state and 0, dissociated state of the ternary complex. The number percentage shown on the bottom right represents the fraction of the important collective basis variables in each PC (See Table 1 for details), (b) Free energy as a function of PC1 and PC3. Representative structures are shown on the surface as indicated by arrows. Note that the minor basin for the binary-Fis configuration is highlighted with a black ellipsoid.

**Table 1.** Coefficients for principal components (PCs)

| | $E_{elec}(1+2)^a$ | $E_{elec}(1)$ | $E_{elec}(2)$ | $N_{contact}^b$ | Dist<sup>c</sup> | $H_{contact}^d$ | $Q_{binFis}^e$ |
| --- | --- | --- | --- | --- | --- | --- | --- |
| PC1 (51.6%)<sup>f</sup> | **0.228<sup>g</sup>** | **0.193** | **0.192** | 0.049 | 0.078 | 0.092 | **0.168** |
| PC2 (20.4%) | 0.107 | 0.086 | 0.094 | **0.368** | **0.210** | 0.118 | 0.017 |
| PC3 (14.0%) | 0.021 | 0.015 | 0.020 | 0.098 | **0.222** | **0.408** | **0.216** |
| PC4 (6.0%) | 0.005 | 0.013 | 0.052 | **0.456** | **0.461** | 0.013 | 0 |
| PC5 (4.5%) | 0.003 | **0.468** | **0.297** | 0.024 | 0.028 | 0.054 | 0.126 |
| PC6 (3.5%) | 0.010 | 0.051 | 0.146 | 0.004 | 0.001 | **0.316** | **0.472** |
| PC7 (0.0%) | **0.628** | **0.174** | **0.198** | 0 | 0 | 0 | 0 |

<sup>a</sup> $E_{elec}(1+2)$ : Sum of electrostatic energy $E_{elec}(\text{Fis1:DNA})$ , between Fis 1 and DNA, and electrostatic energy $E_{elec}(\text{Fis2:DNA})$ , between Fis 2 and DNA. Simple notations $E_{elec}(1)$ and $E_{elec}(2)$ are used to refer to $E_{elec}(\text{Fis1:DNA})$ and $E_{elec}(\text{Fis2:DNA})$ , respectively.
<sup>b</sup> $N_{contact}$ : Total number of contacts between Fis 1 and Fis 2 on DNA. The cutoff for forming a contact ( $C_\alpha$ - $C_\alpha$ ) at the interface is set to be 9.5 Å.
<sup>c</sup> Dist: Distance between the the center-of-mass of Fis 1 and the center-of-mass of Fis 2.
<sup>d</sup> $H_{contact}$ : Specific head-to-head contact between Fis 1 and Fis 2. Only the first 10 residues from the N-terminus of the corresponding chain in the Fis protein are taken into account.
<sup>e</sup> $Q_{binFis}$ : The $Q$ value of binary-Fis configuration.
<sup>f</sup> Percentage of the total variance explained by each PC.
<sup>g</sup> Square of the principal component coefficients shown. This value represents the fraction for the particular collective basis variable in each PC.
Note that $\sum_i^7 f_i = 1$ where i refers to the index for collective basis variable.

## Multiple dissociation pathways

Collective variable (CV) principal component analysis allows one to uncover better dimensions for exploring the population of specific intermediates on the free energy landscape. Fig.4(a) shows the free energy surface as a function of the PC1 and the electrostatic interaction energy *E_elec_*(Fis 1:DNA) and sketches out possible dissociation pathways. In general, the dissociation of one of the proteins Fis1 from DNA progresses as *E_elec_* (Fis1:DNA) increases, and protein Fis1 finally becomes physically separated from DNA in the major basin. During this progression, the electrostatic interaction energy between Fis1 and DNA asymptotically approaches 0, meaning the proteins have become “fully dissociated.” Yet the two Fis proteins can also interact with each other while on the DNA and form binary species. By tracing these populations on the surface, two dissociation pathways are sketched out. The lower arrows (*B* →*I*_1_→*D*_1_) follow the dissociation channel that leads to the net effect of “exchange.” In this exchange process, the second protein molecule Fis2 (red) competes with Fis1 (blue) for the same DNA binding site. As a result, the blue Fis molecule becomes partially dissociated (*I*_1_) and then eventually comes off. The molecular details of this exchange process have been fully discussed in our previous work.^8^ We also find however that dissociation can take place through a distinct route leading to different stoichiometries of reaction, as indicated by the other arrow branching out from *I*_1_, towards *I*_2_ (binary-Fis species) all the way to complete dissociation of both Fis molecules (*D*_2_), namely, “cooperative dissociation.” This alternative dissociation pathway relies on there being an interaction between the two Fis proteins on DNA, in which they form a binary-Fis molecular dyad (*I*_2_) prior to dissociation. We notice that the binary-Fis species manifest themselves as a structural ensemble of unusual protein-DNA ternary assemblies, with multiple configurations dynamically switching between each other along the DNA (see Fig.S3 in Supporting Information). This result suggests that the formation of the symmetric binary-Fis configuration (*I*_2_) on DNA is a prerequisite for carrying out cooperative dissociation. Cooperative dissociation is therefore a biological process that utilizes protein-protein interactions while the proteins are on the DNA.

**Figure 4.**
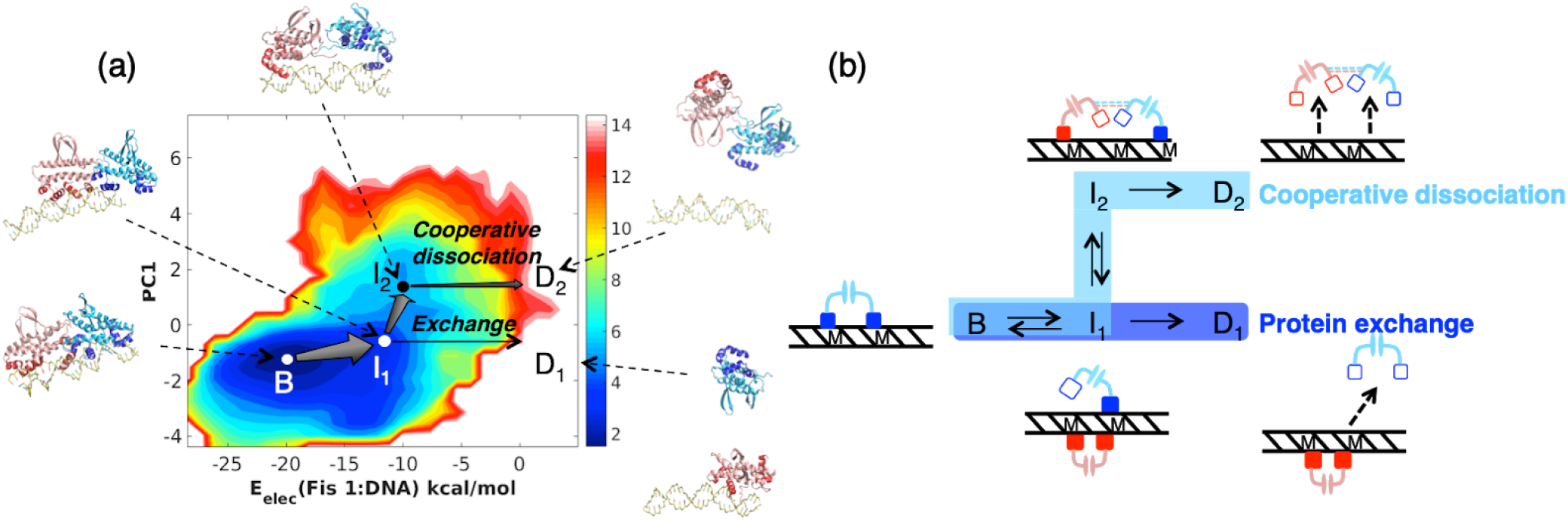
Multiple dissociation pathways of Fis protein from DNA are illustrated, (a) The two-dimensional free energy surface as a function of the electrostatic interaction energy *E*_elec_(Fis1:DNA) and the first principal component PC 1. The free energy surface reveals a population of the metastable binary-Fis configurations, denoted by *I*_2_. *B* refers to the bound state; *I*_1_, refers to the partially dissociated state; *D*_1_ represents one of the dissociated states (the blue Fis is dissociated but the red Fis remains bound.) while *D*_2_ stands for a different dissociated state in which there is a cooperative interaction (both Fis molecules are dissociated). Representative structures are shown along with their two constituent dissociation pathways: 1. *B* → *I*_1_ →*D*_1_, for “facilitated dissociation” and 2. *B* → *I*_1_ → *I*_2_ → *D*_2_ for “cooperative dissociation.” Both pathways are sketched out using kinetic fluxes with arrows pointing out their direction and the width of the arrow head representing the magnitude, (b) Schematic diagram for the two different dissociation pathways. These pathways are color coded: protein exchange (blue) and cooperative dissociation (cyan).

We can compare the kinetic fluxes along these paths based on the assumption that the dissociation occurs through a series of discrete conformational states. One path, *B* →*I*_1_, →*D*_1_ represents facilitated dissociation^8^ via exchange where *B* refers to the bound state, *I*_1_, refers to the partially dissociated state, and *D*_1_, the dissociated state. While *B* →*I*_1_ →*I*_2_ →*D*_2_ describes cooperative dissociation where *I*_2_ denotes a different partially dissociated state in which specific protein-protein contacts have formed with the second Fis while still remaining on DNA; *D*_2_ refers to the other dissociated state that occurs through *I*_2_. These two channels are shown simultaneously on the free energy surface in Fig.4(a). The figure illustrates the two dissociation pathways of the Fis protein from DNA along with their kinetic fluxes. The kinetic flux describes the magnitude and direction of the flow between conformational states, as indicated by arrows (See Methods for details). The width of the arrow heads represents the magnitude of the flux. Comparing the kinetic fluxes between relevant conformational states allow us to assess the significance of each particular pathway from the others. For example, one can use the fluxes pertaining to the distinct dissociation pathways in order to understand of the significance of cooperative dissociation in comparison with facilitated dissociation. As can be seen from the figure, the flux of *B* →*I*_1_ is significantly large (2.89 × 10^5^ (s^−1^); see **Methods** for others). The intermediate *I*_1_, therefore can accumulate rapidly. This rapid accumulation reflects the fact that there is rapid dynamic equilibrium between *B* and *I*_1_. In fact, *I*_1_, plays an important role in modulating different dissociation pathways. Once *I*_1_, forms, it can either proceed to transform directly into *D*_1_, via facilitated dissociation or may undergo a structural rearrangement first into *I*_2_ and then to *D*_2_ through cooperative dissociation. By comparing the flux of the *I*_1_, → *D*_1_), pathway and that for the *I*_1_, → *I*_2_ pathway, we find that the flux of the former pathway (to *D*_1_),) is considerably smaller than that for the alternative pathway (to *I*_2_). This result shows that the transformation of *I*_1_, into *I*_2_ occurs more frequently than the transformation of *I*_1_, into *D*_1_ underlining the significance of there being this alternative dissociation pathway via *I*_2_. A cartoon diagram for describing the two dissociation pathways using these conformational states is schematically shown in Fig.4(b).

### Protein-protein interaction energy and cooperative effects on DNA–a theoretical prediction

The free energy landscape analysis gives much information about the possible conformational states that can be visited along dissociation pathways. The pathways track the way individual chemical species transform into others in a stepwise manner. This pathway analysis allows us also to understand how specific protein-protein interactions influence the cooperative dissociation. Since free energy is a state function, we can construct thermodynamic cycles based on the pathways depicted using chemical equations and then use Hess’s law to disentangle the relevant thermodynamics. Fig.5 presents a reaction scheme that reformulates both facilitated and cooperative dissociation in terms of a thermodynamic cycle, *e.g., <u>B</u>*→*I*_1_→*I*_2_→*D*_2_ →**D**_3_→*D*_1_→*I*_1_→<u>*B*</u> (“_” refers to both the *start* and *end* species of the cycle). The thermodynamic cycle is built by assuming a hypothetical state ***D***_3_ (in bold), where Fis1, Fis2, and DNA are all fully separated from each other. Since ***D***_3_ chemically is an on-pathway species with respect to both *D*_1_ and *D*_2_, this allows us to depict different kinetic pathways from *B* all the way to ***D***_3_. e.g., *B*→*I*_1_ →*D*_1_ →***D***_3_ (which involves facilitated dissociation pathway: *B*→*I*→*D*_1_ and *B*→*I*_1_→*I*_2_→*D*_2_→***D***_3_ (which involves cooperative dissociation pathway: *B*→*I*_1_→*I*_2_→*D*_2_) We notice that the molecular details of the two extended pathways *D*_1_→***D*_3_** (*i*= 1,2) involve two different specific types of interactions. For example, *D*_1_→***D***_3_ involves the dissociation of a single Fis molecule from the DNA (protein-DNA interaction) while *D*_2_→***D***_3_ involves primarily the dissociation of one Fis molecule from the other Fis (see Fig.5 for illustration). The free energy changes involved in each of these processes in principle can be obtained from our free energy analysis. Specifically, the free energies of states *B, I*_1_ *I*_2_,*D*_1_ and *D*_2_ can be directly obtained from the computed free energy landscape. Taking *B* as a free energy point of reference (*F*_B_ =0). the free energy of other states are thus obtained and summarized in Table 2. Using the thermodynamic cycle, one can connect the total free energies added up through the two different pathways

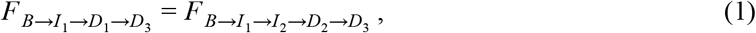

where the left-hand side sums over 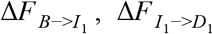 and 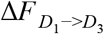; the right-hand side sumes over 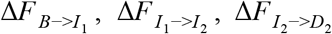, and 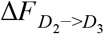. Given that 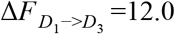 kcal/mol (from our previous work^8^), we obtain from the computed free energy landscape that 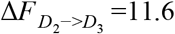 kcal/mol. This value sets an interaction energy scale between proteins to be ~ 10 kcal/mol, an upper bound for the cooperative coupling between Fis proteins. Ideally, the cooperative effect should be estimated considering the presence of the DNA, as can be estimated from conformational state *B* to *I*_2_. This process refers to a structural rearrangement of two bound Fis molecules into the symmetric binary-Fis configuration on DNA through cooperative interaction. The free energy Δ*F*_*B*→*I*_2__ involved in this process gives an approximate value of *J*_coor_ =3.5 kcal/mol, an estimate that is well within the energy upper bound. We also notice that realizing the cooperative interaction in fact involves two mechanistic steps: *B*→I_1_→I_2_. These sequential structural rearrangements occur through the competition for the DNA-binding site. Here we show that our free energy landscape analysis of both dissociation processes provides insight into the cooperative coupling of the proteins while on DNA in a quantitative fashion.

**Figure 5.**
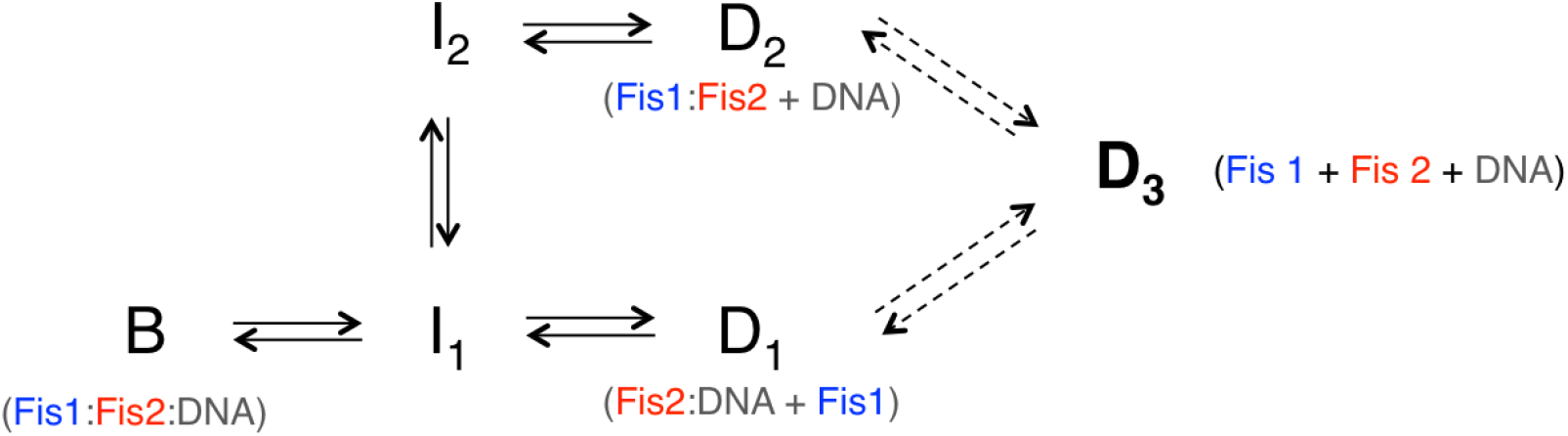
A thermodynamic cycle for the multiple dissociation pathways are shown. The cycle is closed by assuming a hypothetical state *D*_3_, where Fis1, Fis2, and DNA are fully separated, *B* →*I*_1_, →*D*_1_, refers to the pathway for facilitated dissociation while *B* → *I*_1_, → *I*_2_ → *D*_2_ stands for cooperative dissociation. The former leads to protein exchange whereas the latter results in full dissociation of both of the two proteins on DNA. Extended paths *D*_1_, → *D*_3_ and *D*_2_ → *D*_3_ (described in dashed arrows) are used to complete the thermodynamic cycle.

**Table 2.**
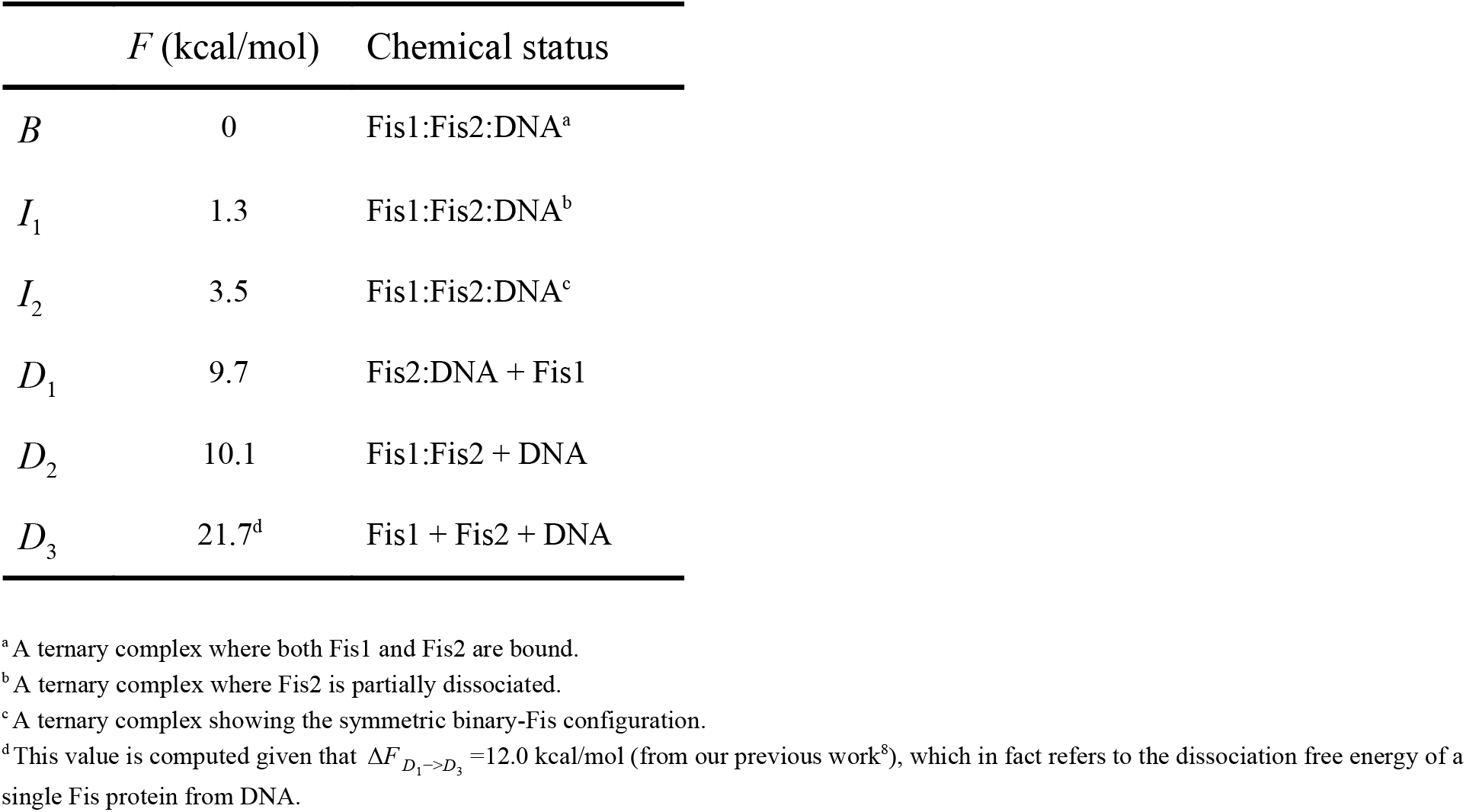
Free energies of conformational states on PC1 free energy surface

### Cooperative dissociation can occur at low protein concentrations regime (~nM)

What is the significance of cooperative dissociation in comparison with facilitated dissociation? Facilitated dissociation refers to a phenomenon of concentration-dependent dissociation of a specific labeled protein molecule from the DNA. As the protein concentration in solution increases, the dissociation rate of labeled molecules also increases. The net effect of facilitated dissociation of a one specific possibly labeled molecule from the DNA however is only that one protein that is already bound on DNA is replaced by another protein of the same type from solution. This is significant for the *in vitro* experimental setup studied by Graham et al.^6^ but would not directly influence the systems biology *in vivo.* Cooperative dissociation, on the other hand, which has a similar concentration dependence by changing the stoichiometry is biologically relevant potentially *in vivo.* It relies on there being an additional structural rearrangement of the two proteins on the DNA which is made possible by protein-protein interaction. To answer the question regarding the significance of cooperative dissociation, or to be precise to, understand what cooperative dissociation would become significant, we need to resort to modeling that describes how the cooperative interactions come into play. A Potts-like statistical model offers a framework to characterize the cooperative binding and unbinding of two proteins. For a single Fis on DNA, we have shown that there are three possible configurations: Bound, partially dissociated, and dissociated states. To describe a combination of two Fis proteins being bound results in a total of nine configurations. If we use a ternary variable for each Fis molecule, the Hamiltonian reads

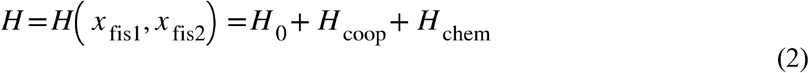

In the equation *H*_0_ refers to the site energy for Fis protein on DNA; *H*_coop_ represents cooperative interaction between the two Fis proteins; *H*_chem_ indicates the chemical potential of the Fis protein; *x*_fis_ is a ternary variable that takes a value of either 0 (dissociated), 1 (partially dissociated), or 2 (bound). See Fig.6(a). Detailed descriptions can be found in the **Methods** section. Fig.6(b) and Fig.6(c) schematically show the configurations of the two Fis proteins on DNA in terms of the ternary variable representation as well as in the dissociation pathways, respectively. The resulting simple statistical model allows one to investigate the effects of cooperative interaction and concentration dependence on the population of binary-Fis configuration. Note that the Hamiltonian shown in Eq. (2) can be generalized to cases where many Fis proteins bind along a long DNA chain, coating the chain, by using a transfer matrix formulation of the partition function,^7^ a task we leave for the future.

**Figure 6.**
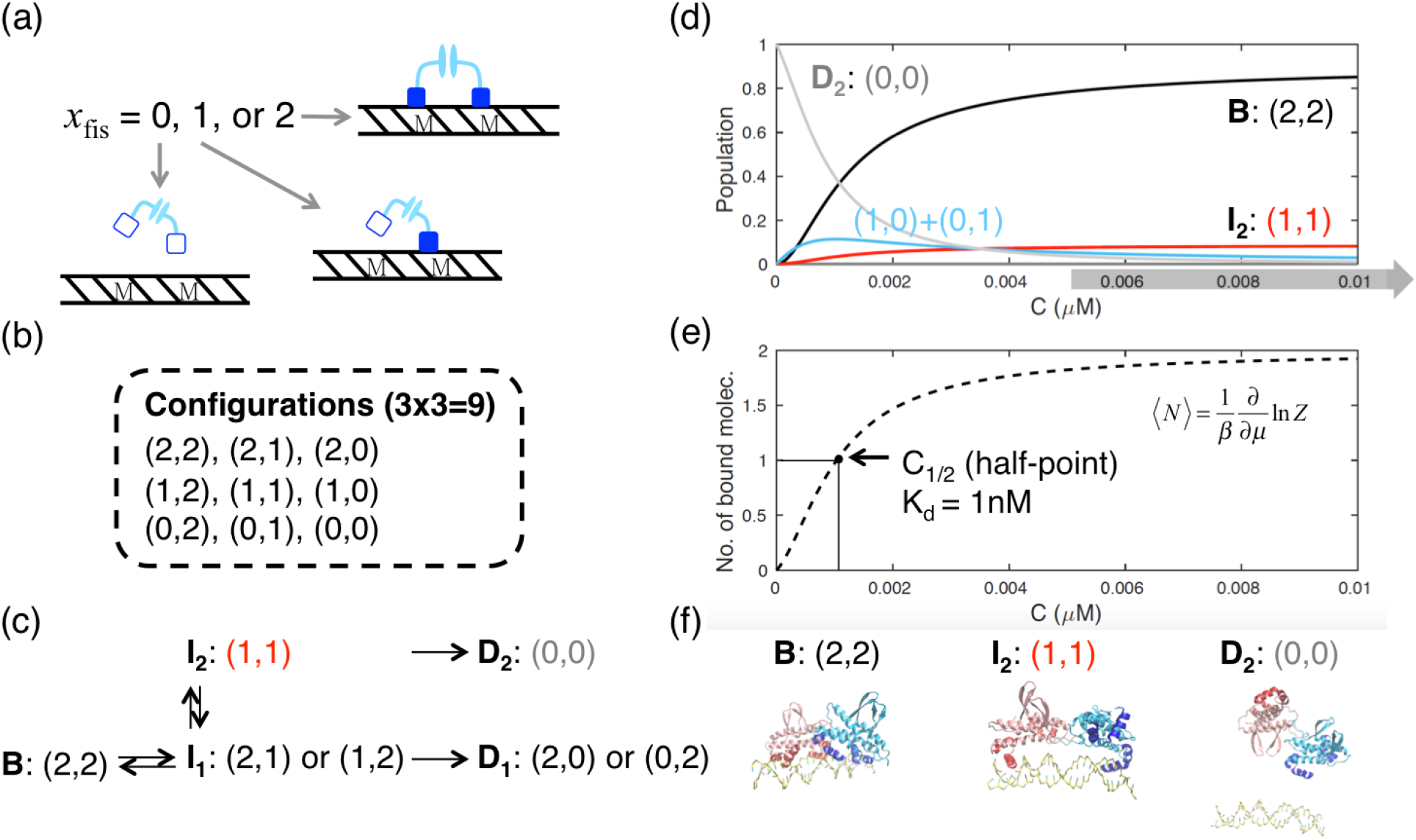
Simple three-state statistical model for describing two Fis proteins binding along DNA is shown, (a) Schematic diagram of the ternary configurations for a single Fis on DNA. (b) Total configurations of two Fis proteins in terms of ternary variables, (c) Binary-Fis species are shown in the dissociation pathways, (d) Equilibrium populations of binary-Fis species are shown as a function of protein concentration C. The gray thick arrow strip shown below indicates the range of concentrations at which the experiments measuring protein dissociation have been carried out i.e., 5~50 nM.^6^ (e) Theoretical equilibrium binding curve with *K*_d_= 1 nM, showing the number of bound molecules (*N*= 2 in total) as a function of protein concentration. This curve is calculated using the 〈*N*=*β*^−1^ ∂lnZ/*∂μ* partition function derivative where *β*=1/*kT* and *μ* is the chemical potential of a single Fis protein, (f) Representative structures for *B*, *I*_2_ and *D*_2_ which are relevant species in the cooperative dissociation. The equilibrium population curves are plotted using 2*ε*_d_=12.0 kcal/mol, *J*=-10.6 kcal/mol, and *μ*_0_ = 5.6 kcal/mol.

Using the simple three-state model describing two Fis proteins binding along the DNA, Fig.6(d) presents the populations of different binary-Fis species as a function of protein concentration. The parameters were obtained and assigned from our free energy landscape calculation. As can be seen from the figure, the population of the bound state *B*, with a configuration of (2,2), increases as the concentration increases. The bound state is the most populated species in the concentration range 5 ~ 50 nM, explored by experiment.^6^ This generic trend captured in the modeling is validated by fitting our theoretical equilibrium binding curve of Fis protein such that the midpoint *C*_1/2_ matches the experimental dissociation constant of the protein *K*_d_ (see Fig.6(e) and **Methods** for details). It is worth noting that the population of the symmetric binary-Fis configuration *I*_2_ also increases with increasing concentration in the experimental concentration regime. This result suggests the importance of the cooperative interactions for permitting cooperative dissociation. Our theoretical modeling supports the idea that cooperative dissociation takes place in the nM concentration regime relevant for most of the transcription factors in the cell.

## Discussion

### Formation of a molecular dyad may play a role in transcriptional activation of genes

In the regulation of genes, there is often an interplay between multiple biological components/regulators binding to the same target site or to non-coding control sequences. In some cases, protein regulators may form clusters on the sequence. Presumably, these protein clusters are able to exhibit multiple structural configurations on DNA so they carry out distinct regulatory tasks in response to different cellular circumstances. We have found that Fis proteins manifest multiple configurations by forming a molecular dyad. This binary-Fis intermediate is formed via a linkage of contacts between two N-terminal coiled peptide segments from the individual Fis proteins. The formation of this molecular dyad allows multiple alternative dissociation pathways, *i.e.,* both facilitated dissociation of a single Fis via exchange and cooperative dissociation of pairs of Fis molecules. This cooperative dissociation differs from the facilitated dissociation studied in the single molecule imaging experiments in terms of the stoichiometry of the dissociation. Biologically, these two paths reflect the different choices that can be made when cells respond to different external stimuli. It is also important to point out the distinct role of the N-terminal coiled region of a single Fis in combining two Fis proteins into one Fis structural dyad. This protein dyad turns out to be an essential element for mediating the different dissociation pathways. The discovery of the binary-Fis dyad reveals a potential functional role of the protein dyad as a molecular switch in dissociation (see below).

Fis protein is known to function as a global transcription factor that can positively or negatively regulate transcription of many genes in bacteria.^5^ For example, Fis can activate transcription when binding to the promoter region that overlaps the binding site of RNA polymerase (RNAP).^22,23^ Many studies have shown that transcriptional activation of genes is mediated by direct interaction between specific amino acids within a turn connecting the B and C helices in Fis (the BC turn) and the C-terminal domain of the α-subunit of RNA polymerase (α-CTD of RNAP). Cheng et al. found that the residues localizing to the BC turn of Fis form a ridge that could contact the α-CTD of RNAP on one side.^24^ These residues include Gln68, Arg71, Gly72, and Gln74. Since the formation of the contacts between the Fis BC turn and the RNAP α-CTD unit requires a specific orientation of the dimerized Fis proteins with respect to the DNA, this structural constraint very likely helps determine the probability of forming the binary-Fis molecular dyad. In other words, the binary-Fis molecular dyad plays a role in modulating Fis’s transcriptional activation by either interrupting or enhancing Fis’s contact with the RNAP α-CTD unit. To test this, we have carried out a simple free energy perturbation survey over several Fis mutants whose effect on transcriptional activation has been reported.^24^ We examine the predicted changes of the population of the binary-Fis molecular dyad on the free energy landscape for these different Fis mutants (see Fig.S4) and compare the results with their corresponding transcription activity *in vivo.* The transcription activation activity in the laboratory was reported at three different levels: high (wild-type), low (20-40 % of wild-type activity), and very low (<10 % of wild-type activity). We see from our perturbed free energy surfaces that the stability of the binary-Fis molecular dyad with respect to the bound state for a range of mutants seems to exhibit some correlation with their *in vivo* transcription activation activity (see Fig.S5), suggesting a possible role for the binary-Fis molecular dyad in modulating the process of the transcription activation.

Depending on the types of transcriptional activators (class I or II), the extent of the interference may vary. For the *rrnB* P1 promotor, Fis functions as a class I transcriptional activator binding upstream of the binding site for the σ-70 form of RNAP.^23^ For the *proP* P2 promotor, Fis functions as a class II transcriptional activator binding at the site overlapping the −35 binding site of the σ-38 form of RNAP.^25^ Because class I and II activators involve different binding sites and different types of transcription assemblies formed, transcription initiation may involve distinct binary-Fis populations at the same overall Fis expression level. The extent of transcriptional activation therefore can be different. In other words, a group of functionally related genes, given that they are regulated via similar mechanisms, can be controlled all at once through the modulation of protein expression level. This study shows that multiple binding configurations of transcription factors are the key to uncovering structure-function relationships for transcriptional regulation by Fis. The mechanism described may be the prototype for explaining the growth phase-dependent gene regulation profile observed from experiments.^5^

### Molecular mechanisms underlying cooperative dissociation and its relation to molecular stripping

Unlike a conventional genetic switch, which relies solely on protein-DNA binding affinities under equilibrium,^26^ kinetically controlled dissociation leads to a non-equilibrium response to external stimuli. The residence time of a transcription factor on DNA binding sites is concentration dependent. Our previous work showed that the concentration dependence of dissociation arises from the formation of a ternary intermediate (Fis1:Fis2:DNA) where the originally bound protein Fis1 becomes partially dissociated due to the solution protein Fis2 competing (with Fis1) for the DNA binding site.^8^ In other words, the phenomenon of facilitated dissociation originates from DNA binding site competition.

A somewhat analogous situation where one kind of protein molecule catalyzes the dissociation of a transcription factor from DNA has been reported experimentally and analyzed computationally for the *NFκB/Iκ*B/DNA regulation network.^27–30^ The molecular mechanism in that case however is fundamentally different from the one described here. The dissociation rate of *NFκB* from DNA increases as the solution-phase concentration of *IκB* increases, not as the concentration of *NFκB* itself increases, although the latter is potentially a possibility that has not yet been explored in the laboratory.^28^ The catalyzed dissociation process involves a functional domain twist mode activated by the binding of *IκB* to *NFκB* in the ternary complex (with DNA).^27^ In this case, the stripping molecule *IκB* directly interacts with *NFκB* through protein-protein interactions and then removes *NFκB* from DNA by lowering its binding affinity with DNA.^27,30^ In contrast, facilitated dissociation of Fis proteins arises from DNA binding site competition through protein-DNA interactions, a strategy that may also be utilized in situation like chromosome packaging.

Cooperative dissociation is a complex process that involves both protein-protein and protein-DNA interactions at different stages. In the first stage, an incoming protein molecule Fis2 binds to the DNA in the Fis1-DNA complex through non-specific protein-DNA interactions. Because of the protein invasion, the resident protein molecule Fis1 becomes partially dissociated.^8^ In the second stage, the two Fis molecules on DNA undergo an orientational rearrangement into a molecular dyad via protein-protein interactions. Following this event, the binary-Fis molecular dyad can dissociate as a whole. Consequently, cooperative dissociation consists of sequential events involving both protein-protein and protein-DNA interactions. In summary, cooperative dissociation is initiated by protein-DNA interaction first which is then followed by protein-protein interaction prior to separation from DNA.

### Potential role of cooperative dissociation in the systems biology of gene regulation by Fis

Although single-molecule experiments have revealed the anomalous dissociation behavior of Fis molecules from DNA *(i.e.,* facilitated dissociation), facilitated dissociation results in the net effect of “exchange of protein”. Due to the unchanged stoichiometry of the [], facilitated dissociation presumably does not influence the gene regulation by Fis at the systems-level. As a master transcription regulator, Fis protein can regulate over two hundred genes of *E-coli*.^5^ Classical models suggest that the thermodynamics of binding and unbinding of proteins on DNA acts as the molecular switch for activating or deactivating a gene. This two-state thermodynamic model unequivocally attributes the activation of gene expression to the binding of a Fis molecule. Facilitated dissociation mainly causes replacement of Fis molecule already bound on DNA by a solution-phase one. The overall Fis regulatory function would not be influenced by such a replacement event, though this process is still under kinetic controlled. In contrast, the process of cooperative dissociation presented in this work not only reaffirms the significance of kinetic controlled protein dissociation from DNA, but also emphasizes the role of the binary-Fis molecular dyad in the systems biology of the Fis regulatory network. The formation of the cooperative binary-Fis configuration prior to dissociation (of both Fis molecules from DNA) is analogous to the intermediate process of molecular stripping of *NFκB/Iκ*B/DNA regulation network where the formation of the *NFκB-Iκ*B co-complex on DNA represents a transient ternary structure formed before dissociating from DNA. Since the stripping process releases all the *NFκB* molecules at once and quenches transcription, the systems biology of this *NFκB/Iκ*B/DNA regulation network is very likely also relevant for the systems biology of gene regulation by Fis. We are now investigating Fis-related transcription components. This type of survey will allow us clarify their cross-correlation behavior within the Fis regulation network at the systems-level. Notably, it has been reported that facilitated dissociation of Fis can also be carried out by other proteins such as HU and NHP6A.^6^ Facilitated dissociation can be rather generic among a variety of transcription factors.

### A suggestion for experimentally verifying the existence of the binary-Fis molecular dyad and of multiple dissociation pathways

Our theoretical predictions for cooperative dissociation open a door to mechanistic understanding of transcriptional regulation by Fis. We notice that the publishing kinetic studies on Fis in the laboratory, however, are not able to distinguish the exchange process from the cooperative dissociation process.^6^ Accordingly, we propose an experimental scheme to distinguish the pathways and detect the presence of the binary-Fis intermediate and its subsequent dissociation. Previously, Chen and coworkers used a single-molecule FRET technique along with surface immobilization of DNA to study the structural dynamics of a metalloregulator CueR on DNA.^21^ Three conformational states of CueR on DNA were distinguished using single-molecule *E*_FRET_ trajectory of an immobilized Cy3-DNA interacting with Cy5 labeled CueR. These three states include one free DNA state and two binding orientations of the labeled CueR on DNA. In addition, they used an engineered DNA Holliday junction (HJ) along with the same FRET technique to probe the interaction between multiple CueR molecules and the engineered DNA. They specifically observed a biphasic protein concentration dependence of the time-averaged interconversion rate between the two HJ conformers (conf-II → conf-I), an indication of forming a CueR1:CueR2:DNA ternary complex. Similar single-molecule FRET techniques can be applied to study the conformational dynamics of Fis on DNA, in particular, to probe termolecular interaction that at some time point lead to the formation of a transient binary-Fis ternary complex (Fis1:Fis2:DNA). Since our simulations reveal an ensemble of configurations for Fis protein pairs, it would be interesting to see from experiments whether the interconversion rates between them show biphasic or multiphasic protein concentration dependence. Meanwhile, clustering analysis may be applied, along with the maximum-information method taking into account photon-counting statistics,^31^ to interpreting the FRET experiment proposed in order to distinguish different binary-Fis species from others in a statistically robust manner.

To distinguish dissociation pathways, we suggest the use of multiple fluorescence probes. In the single-molecule experiment tracking protein dissociation from DNA, two stages are involved. In the first stage, fluorescence dye labeled protein is introduced to a flow cell where an extended DNA is tethered to the glass surface with magnetic tweezers. The DNA is thus coated by protein molecules. The dissociation of the protein from DNA can be monitored by recording the change of fluorescence intensity on DNA using fluorescence microscopy. In the second stage, one can use protein labeled with a different fluorescence dye that forms a donor-acceptor pair of FRET. In addition to tracking the fluorescence intensity change coming from the coated protein molecules, one can also specifically track the presence of the incoming protein molecules from solution and correlate their dissociation behavior from DNA with the formation of intermediates using FRET efficiency.

## Conclusions

In this study, we carry out molecular dynamics simulation combining the state-of-the-art coarse-grained protein force field (AWSEM) with a state-of-the-art coarse-grained DNA force field (3SPN.2c) to explore the global binding energy landscape of protein-DNA assemblies. Where two Fis proteins interact with each other on the DNA, simulation uncovers a distinct binary-Fis molecular dyad whose formation relies on complex structural rearrangement of the two molecules. We find that this binary-Fis configuration shows a C_2_ chemical symmetry on the DNA that overall represents a unique ternary intermediate prior to cooperative dissociation. This cooperative effect occurs between the two proteins via protein-protein interaction. The collective variable principal component analysis that we developed shows the cooperative interaction between the two Fis molecules on the DNA can eventually lead to co-dissociation of both molecules simultaneously. The multiple protein dissociation pathways that we discovered have important implication for understanding protein’s stoichiometries upon dissociation. In particular, our findings support the idea that the capability of forming multiple binding configurations on DNA allows the proteins to modulate their own dissociation pathways so as to carry out distinct regulatory tasks in response to different cellular circumstances. When protein dissociates through “facilitated dissociation,” the protein on the DNA is replaced by solution-phase protein through competition for DNA binding sites. The net effect of this process is “protein exchange.” In the process of gene expression, transcription factors in action very often require constant renewal in order to maintain the robustness of their regulatory tasks, in particular for genes that switch between different chromosome packaging states. On the other hand, when a solution-phase incoming protein interacts with the protein on DNA, they can form multiple binding configurations through the cooperative interaction. Co-dissociation therefore may occur once the binary-Fis molecular dyad is formed, a process akin to the “molecular stripping” seen in the NFκB/IκB genetic broadcasting system. To quantitatively understand how the cooperative effect influences protein dissociation from DNA, we use a three-state binding model that incorporates several binding parameters to study the population of multiple binding configurations in response to varying protein concentration. We find that the energy scale for the cooperative interaction is roughly <10 kcal/mol. This value ensures the meta-stability of the binary-Fis molecular dyad in ~nM protein concentration, a typical range within cells. Our simulation results provide kinetic insights into the systems biology of gene regulation by Fis.

## Methods

### Coarse-grained models for proteins and DNAs

The simulations in the present study were carried out using a hybrid simulation scheme using AWSEM/3SPN.2C based coarse-grained models for both the proteins and the DNA. For the coarse-grained model of proteins, we used AWSEM (**A**ssociative-memory, **W**ater-mediated, **S**tructure and **E**nergy **M**odel^32^), a model where each amino acid residue is represented by three beads (O, C_α_, and C_β_) while the DNA is modeled using the 3SPN.2C force field.^33,34^ The 3SPN.2C model employs a similar coarse-grained architecture of 3 beads per functional unit (i.e., nucleotide/amino acid residue for DNA/protein). This hybrid AWSEM/3SPN.2C scheme has already been used for a variety of protein-DNA systems.^8,27,35^ The details of the force field models can be found elsewhere.^8^

### *Q* value of binary-Fis configuration (*Q_binFis_*)

A centroid structure taken from the ensemble of binary-Fis configurations was used as a reference structure. This binary-Fis ensemble contains a total of 1100 continuous frames that were selected from the sampling window *E*_elec_= −15 kcal/mol. To measure the structural similarity of snapshots from the trajectories with respect to the binary-Fis reference structure, we used an order parameter *Q*, which compares the pairwise distances of *C*_α_ atoms among the residues in a given instantaneous structure to those in the binary-Fis reference structure. The *Q* value (*Q_binFis_*) for all snapshots from the simulation trajectories was calculated using the following expression

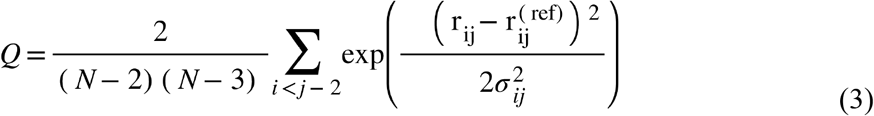

where *N* is the total number of residues; *r_ij_* is the instantaneous distance between *C_α_* atoms of residue *i* and *j* in the Fis protein pair; 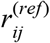 refers to the corresponding distance in the reference structure; *σ_ij_* denotes an accuracy threshold *σ*_ij_=(1+|*i-j*|)^0.15^. Note that the summation in Eq. (3) involves only the pairs that are separated by three or more residues in sequence.

### Collective Variable Principal Component Analysis

To understand the free energy surfaces of the binary-Fis configuration on DNA (Fis 1:Fis2:DNA), we studied the essential dynamics of the system by performing principal component analysis (PCA) on simulation trajectories. Principal component analysis is used to extract essential dynamics as a function of a set of basis variables, e.g., cartesian coordinates. The method generates a new set of variables (principal components) that govern the behavior of the system and that each principal component is a linear combination of the original basis variables. Very often the basis variables are the cartesian coordinates of the atoms or residues but other physically meaningful basis variables have been used such as contact occupations^36,37^ and local strain variables.^38^ In this study, we choose to use a novel type of PCA where the basis variables are chosen from a set of preconceived intuitive global progress coordinates that had previously proven useful in describing collective motions of biomolecules. For instance, in our previous work electrostatic energy (*E*_elec_) between Fis protein and DNA was used to study dissociation mechanism of Fis protein from DNA with great success.^8^ Other physically motivated variables include the total number of protein contacts (*N*_contact_). center-of-mass (COM) distance, head-to-head protein contacts, *Q* value of the binary-Fis configuration. The basis variables adopted in the present study along with their coefficients and percentage of total variance are listed in Table 1.

In this type of PCA, the covariance matrix is built using the *n*-by-*p* simulation data matrix with *n* (rows) corresponding to the number of observations and *p* (columns), the number of collective basis variables. This type of analysis is termed collective variable PCA and was carried out using standard PCA function implemented in Matlab 2015a. By default, PCA centers the data and uses the singular value decomposition (SVD) algorithm. When performing collective variable PCA, we used the inverse of variances of the ingredients as variable weights. A weighted PCA can avoid large errors due to outliers in the data, thus increasing the robustness of the analysis.

### Kinetic flux calculation

The calculation of kinetic flux is based on Kramers theory as described in the supporting information of our previous work.^8^ The Kramers barrier crossing rate is used describe the transition rate between two local minima separated by a barrier on the free energy landscape. The barrier height along with other parameters of the free energy profile can be quantified by projecting free energy onto one specific reaction coordinate. In the overdamped limit, it follows that the thermal escape rate from 1D potential reads

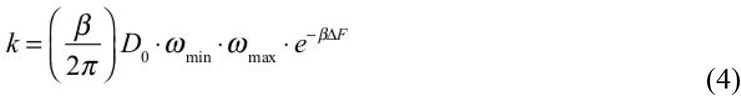

where *β*=1/*k_B_T, D*_0_ refers to the effective diffusion constant on a equivalent flat landscape of the initial free energy minimum basin, *ω*_min_ and *ω*_max_ are the curvature at the minimum and the curvature at the barrier maximum, respectively.

Previously, we have determined *D*_0_ = 6 x 10^7^ (kcal/mol)^2^S^−1^ using the linear relation of mean-squared displacement (MSD) along the reaction coordinate, 〈Δ*x*^2^s_−1_ = 2*D*_0_, ·*t* (*x* is the reaction coordinate chosen). Similarly, *ω*_min_ = *ω*_mm_ = 0.4 (kcal/mol)_-½_ was used. Δ*F* refers to the free energy difference between any two states on the free energy landscape. This free energy difference along with *D*_0_, *ω*_min_, and *ω*_max_ together were used to estimate the relevant kinetic flux. Since *D*_0_, *ω*_min_, and *ω*_max_ are constants, the kinetic fluxes between different states are solely determined by their relative free energy difference. The kinetic fluxes computed were *k*_*B*→*I*_1__ = 2.89 × 10^5^ (s^−1^), *k*_*I*_1_→*D*_1__ = 1.93 (s^−1^), *k*_*I*_1_→*I*_2__ = 6.4 × 10^4^ (s^−1^), *k*_*I*_2_→*D*_2__ = 39.7 (s^−1^) are represented by the width of the arrows and are shown on Fig.4(a). Table 2 shows the free energies of conformational states used in the flux calculation. These free energies are shown with respect to a reference state on the PC1-E_elec_(Fis1:DNA) surface.

### Three-state binding model

The Hamiltonian of the statistical model uses a 3-state Potts-like description. The model includes three terms that represent the site energy of Fis on DNA, cooperative interaction between Fis molecules, and their chemical potential, respectively,

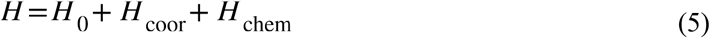

wheri

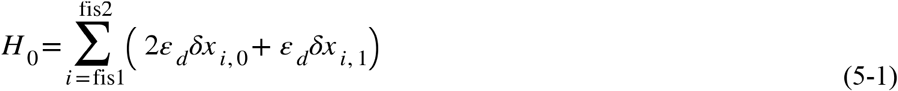

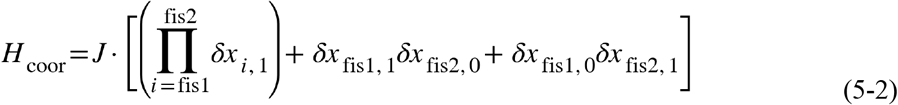

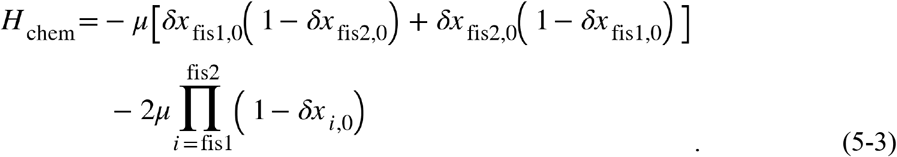

The site energy *H*_0_ describes the free energy change in terms of the dissociation energy *ε_d_* of individual DNA-binding domains. One Fis molecule has two DNA-binding domains. The index *i* in the summation loops over 2 Fis proteins (fis1 and fis2). The kronecker delta *δx_i,j_* denotes

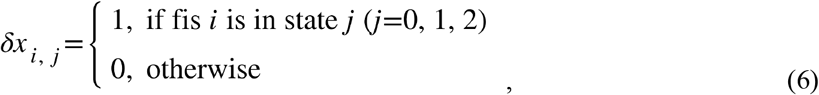

where *j* = 0, 1, 2 refer to the fully dissociated, the partially dissociated, and the bound states, respectively. The first energy term in *H*_0_ describes the configuration of Fis in the dissociated state (*j*=0); the second energy term refers to the configuration of Fis in the partially dissociated state (*j*= 1); the third term is ignored. The cooperative interaction *H*_coor_ describes the correlation between two Fis proteins when on the DNA. *J* is an interaction energy that quantifies the cooperative coupling of binding the two Fis proteins. There are three scenarios for an effective interaction to take place. They are described in three terms. The first term is effective only if both Fis molecules are in the partially dissociated state–a scenario of forming the unique binary-Fis configuration on DNA. The second term describes another scenario of forming the binary-Fis configuration but with only Fis1 in the partially dissociated state on DNA while Fis2 is dissociated. The third term denotes a similar scenario but now with Fis2 being in the partially dissociated state while Fis1 is dissociated. Note that the second and the third terms are used to describe partially dissociated configurations of the molecular dyad on DNA. The last energy component *H*_chem_ takes into account the chemical potential of Fis molecule in solution. The binding of each Fis molecule to DNA contributes -*μ* energy to the Hamiltonian. If both Fis molecules are bound, the system would gain *-2μ* as a result. Note that for Fis molecule being in either state 1 (partially bound) or 2 (fully bound) the Fis in general is considered “bound” on DNA (gaining -*μ*).

## Acknowledgments

The project described was funded by the NSF sponsored Center for Theoretical Biological Physics (Grants PHY-1308264 and PHY-1427654) with additional support from NIH Grant R01 GM44557. Support from the National Institute of General Medical Sciences PPG Grant P01 GM071862 and from the D. R. Bullard-Welch Chair (Grant C-0016) at Rice University to P.G.W. is greatly appreciated.

