## Supporting information for "Multiple Binding Configurations of Fis Protein Pairs on DNA: Facilitated Dissociation versus Cooperative Dissociation"

### Table of Contents

**Figure S1. Free energy surfaces as a function of several of trial collective coordinates are shown. 2**

**Figure S2. The trial collective basis variables that are used in the principal component analysis. 3**

**Figure S3. The structural ensemble of the binary-Fis configuration and their mutual dynamic switching are schematically shown. 4**

**Figure S4. A free energy surface survey (free energy projected onto PC3-PC1) over a range of Fis mutants for the stability change of the binary-Fis molecular dyad is shown. 5**

**Figure S5. The correlation plot of the Fis transcription activity (due to mutation effects) in relation to different thermodynamic properties, obtained from both experiment and simulation, is shown. 6**

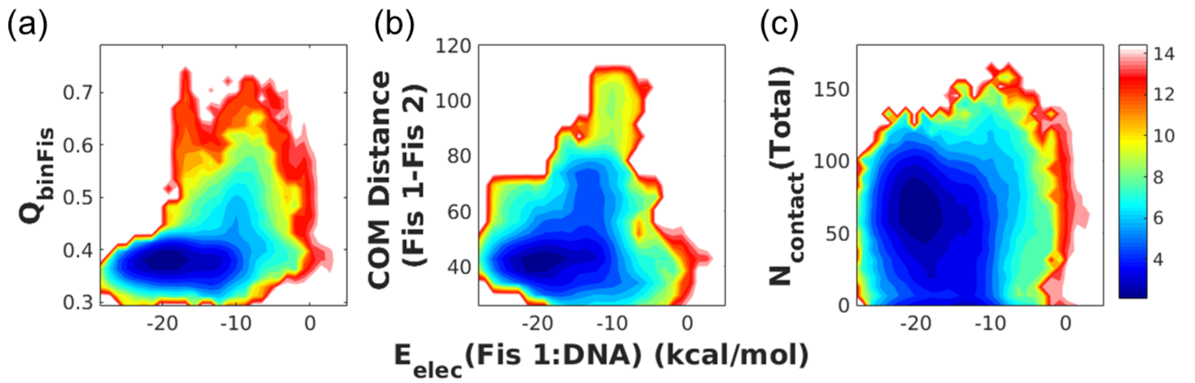


Figure S1. Free energy surfaces as a function of several of trial collective coordinates are shown. (a) $Q$ value of the binary-Fis configuration ($Q_{binFis}$) (b) Center-of-mass distance between Fis1 and Fis2 on the DNA (c) The total number of protein contacts ($N_{contact}$).

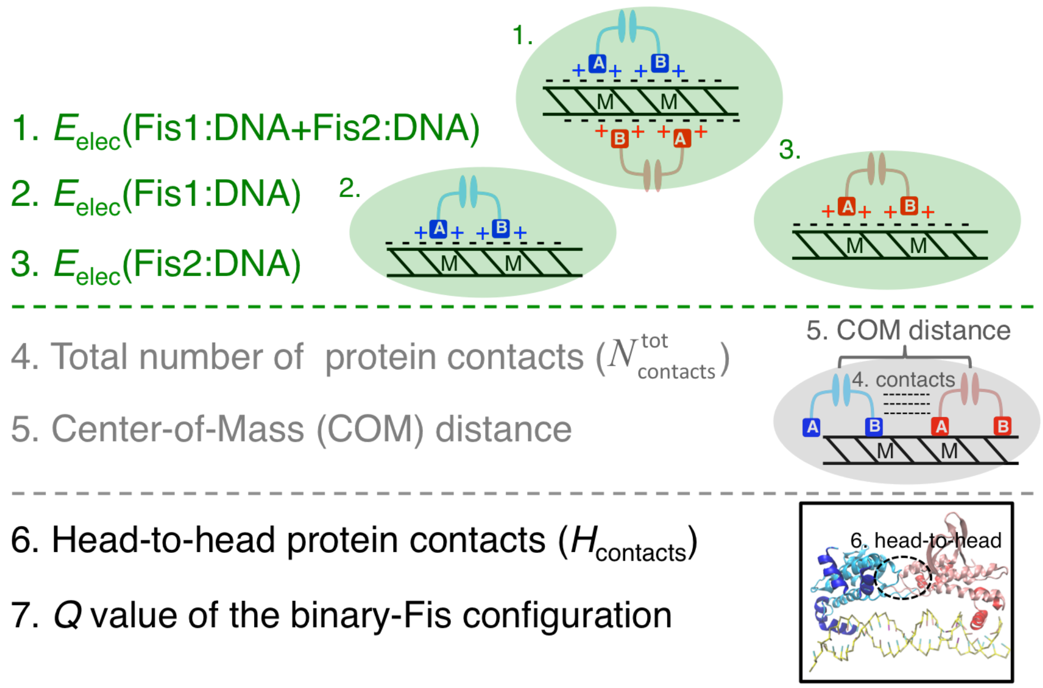


Figure S2. The trial collective basis variables that are used in the principal component analysis. These collective variables are grouped. (*Top*) Items 1-3 (in green) describe the electrostatic interactions between the protein and DNA. (*Middle*) Items 4-5 (in gray) describe physical protein contacts and geometric position between proteins on DNA. (*Bottom*) Items 6-7 describe structural specificity that distinguishes several of the particular protein quaternary structures from each other.

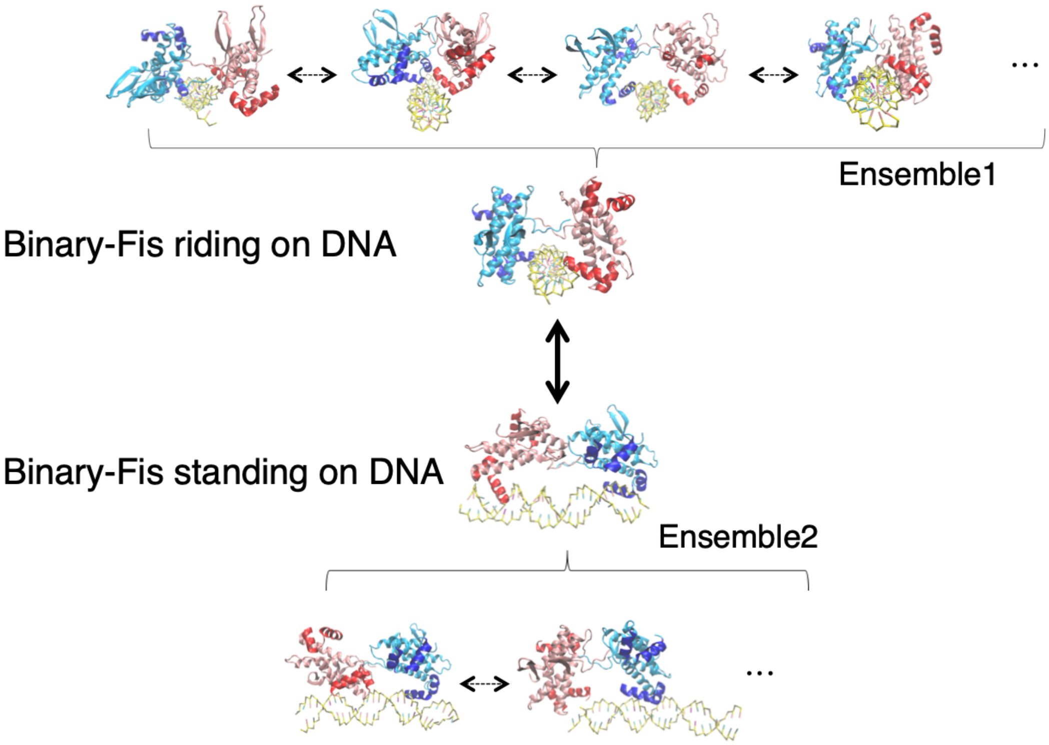


Figure S3. The structural ensemble of the binary-Fis configuration and their mutual dynamic switching are schematically shown.

The structural ensemble of the binary-Fis configuration on DNA contains two major categories of orientation: riding and standing. The binary-Fis structure can dynamically switch between the two distinct orientations by sliding along the DNA, denoted by a *black-thick* line with double arrows. In each of the categories, the structure of the protein-DNA ternary assemblies also shows multiple configurations. The *dashed-thin* lines with double arrows within individual ensemble 1 and 2 describe dynamic switching between each other. Note that only some example structures are shown.

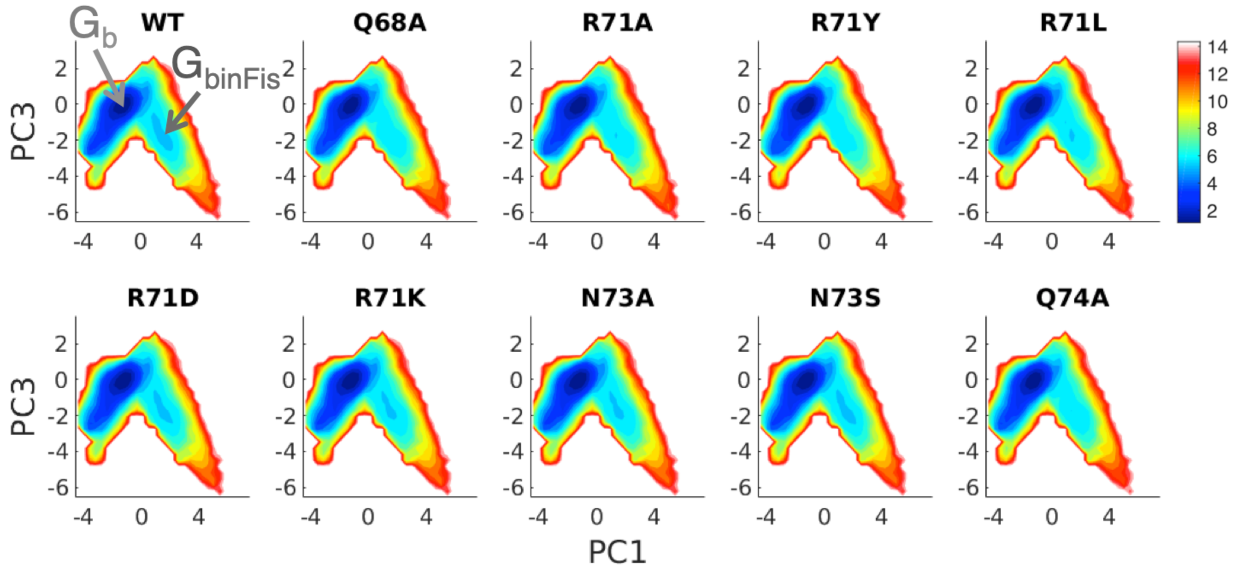


Figure S4. A free energy surface survey (free energy projected onto PC3-PC1) over a range of Fis mutants for the stability change of the binary-Fis molecular dyad is shown.

The free energy of the binary-Fis molecular dyad ($G_{binFis}$, shown in dark gray) is calculated with respect to the bound state ($G_b$, shown in light gray). One can thus calculate the free energy change $\Delta G_{WT} = G_{binFis} - G_b$; Similarly, $\Delta G_{binFis}(mut) = G_{binFis}(mut) - G_b$.

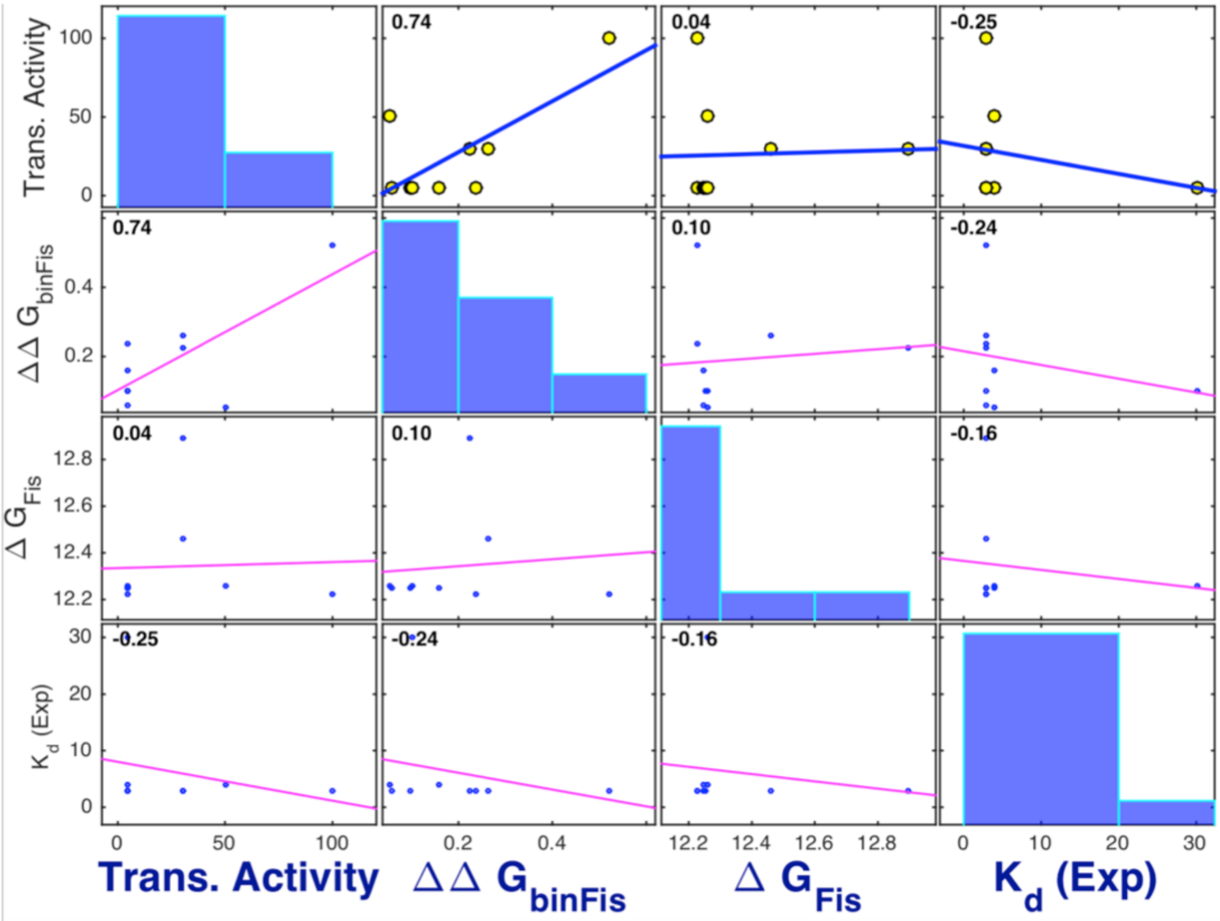


Figure S5. The correlation plot of the Fis transcription activity (due to mutation effects) in relation to different thermodynamic properties, obtained from both experiment and simulation, is shown. Several of the thermodynamic properties such as $\Delta G_{Fis}$(mut), $\Delta\Delta G_{binFis}$(mut), and experimental $K_d$ (dissociation constant[1]) of the mutants are cross-correlated along with their corresponding transcription activity and are compared on the plot. Note that $\Delta\Delta G_{binFis}(mut) = \Delta G_{binFis}(mut) - \Delta G_{WT}$. The number shown on the *top-left* of each panel represents the correlation coefficient. The Fis transcription activity shows a significant correlation with $\Delta\Delta G_{binFis}$ with $\rho_{X,Y}$=0.74. $\Delta G_{Fis}$ is the simulated free energy of dissociation of Fis from DNA.
